# Design and optimisation of rapid real-time PCR assays for the detection of key *Culicoides* species

**DOI:** 10.1101/2025.09.26.678746

**Authors:** Elsie Isiye, Angela Valcarcel Olmeda, Thomas Curran, David O’Neill, Theo De Waal, Gerald Barry, Aidan O’Hanlon, James O’Shaughnessy, Marion England, Annetta Zintl, Denise O’Meara

## Abstract

In extensive surveillance programmes of *Culicoides* biting midges (Diptera: Ceratopogonidae), morphological identification can be time-consuming and difficult, while DNA barcoding, although highly accurate, may not be cost-effective or suitable for rapid analysis, as it requires individual specimen processing. To address these limitations, we developed a rapid screening method using real-time PCR assays with either SYBR Green or hydrolysis probe-based detection chemistries. Species-specific primers and, where necessary, hydrolysis probes were designed based on the updated sequences of the ITS2 region of seven *Culicoides* species. The specificity and efficiency of these assays were validated both *in silico* and through real-time PCR testing on target and non-target *Culicoides* species, tested individually and as mixed-species samples. The new real-time PCR assays detect vector species including *C. obsoletus, C. scoticus, C. chiopterus, C. dewulfi, C. pulicaris, C. punctatus*, and *C. impunctatus* in pools of individual specimens, with single-specimen sensitivity. The molecular techniques developed in this study provide a valuable tool for accurate and high-throughput *Culicoides* surveillance, which can be used for year-round monitoring of adult midges in traps and larvae in environmental samples, potentially revealing novel insights into the spatial and temporal turnover of *Culicoides* species. These methods can be applied to large-scale vector screening programmes involving pooled samples, addressing the limitations of previously described methods used in midge surveillance.

## Introduction

Molecular-based species identification is often a crucial step in ecological surveillance programmes, whether for biodiversity monitoring (Curran et al. 2022), surveillance of protected species (Harrington et al. 2019), invasive species (Browett et al. 2020), or species of interest from a One Health perspective, such as vectors of disease (Isiye et al. 2025). Novel real-time PCR assays have been developed and applied across a range of ecological contexts in Ireland and Britain. Applications include species and sex identification of Eurasian otters (*Lutra lutra*) from non-invasive samples (O’Neill et al. 2013), large-scale monitoring of native bat species (Harrington et al. 2019), and discrimination between native red squirrels (*Sciurus vulgaris*) and invasive grey squirrels (*Sciurus carolinensis*) (O’Meara et al. 2012). Real-time PCR has also been applied to detect target species from indirect sources of DNA, such as faecal or environmental DNA (eDNA) sources (O’Meara et al. 2014). The versatility and accuracy of real-time PCR detection methods can be useful in various ecological surveillance contexts, including vector surveillance.

In Europe, *Culicoides* biting midges are well recognised as both disease vectors and nuisance pests across various regions (Carpenter et al. 2013; Prudhomme et al. 2025). In Ireland, the majority of commonly occurring *Culicoides* species are of significant veterinary importance, as they are well-known vectors of bluetongue virus (BTV) and Schmallenberg virus (SBV) (McCarthy et al. 2009; Collins et al. 2018; Jess et al. 2018; Isiye et al. 2025). The identification of *Culicoides* species has traditionally relied on morphological characteristics, including wing patterns, head structures, and genitalia (Mathieu et al. 2012), a method that has served as the foundation for the identification and classification of insect species. In recent years, DNA barcoding of mitochondrial and ribosomal gene regions has been increasingly used, providing high accuracy in distinguishing closely related species (Mignotte et al. 2020) and identifying new species (Nielsen et al. 2015; Sarvašová et al. 2017). In large-scale vector surveillance programs, both these methods are too labour-intensive and costly, as they require processing of very high numbers of individual specimens and significant expertise to identify the exact species within the sample collection. On the other hand, DNA metabarcoding, although capable of detecting a broad range of species in complex mixtures or pooled samples (Browett et al. 2021; Curran et al. 2022), often identifies numerous non-target organisms and depends on extensive reference databases, which are still limited for groups such as *Culicoides* midges. While genus-specific primers have been successfully used to distinguish *Culicoides* from other Ceratopogonidae (Cêtre-Sossah et al. 2004), species-specific tests would provide a robust approach for simplifying their identification and allowing for targeted surveillance.

The first conventional species-specific PCR tests to be developed targeted *Culicoides imicola* Kieffer, 1913, as it was considered to be the primary vector of BTV and African horse sickness virus (AHSV) in southern Europe and endemic regions of Africa (Mellor & Boorman, 1995; Miranda et al. 2003; Carpenter et al. 2009; Jacquet et al. 2015; Leta et al. 2019). This involved the design of species-specific primers to develop multiplex PCR assays for the detection of C. *imicola* in insect pools (Cêtre-Sossah et al. 2004; Casati et al. 2009). However, as BTV spread further north and SBV emerged, beyond the range of C. *imicola*, additional competent vector species within the subgenera Avaritia and *Culicoides* were identified, including *Culicoides* obsoletus (Meigen, 1818), *Culicoides* scoticus Downes & Kettle, 1952, *Culicoides* dewulfi Goetghebuer, 1936, *Culicoides* chiopterus (Meigen, 1830), *Culicoides* pulicaris, and *Culicoides* punctatus (Meigen, 1804) (Nolan et al. 2007; Gloster et al. 2008; Carpenter et al. 2009; Wilson & Mellor, 2009). This expansion has led to the development of conventional diagnostic PCR tests for the detection of these vector species. Species-specific multiplex PCR assays have been developed for *Culicoides* species identification, utilising both cytochrome c oxidase subunit 1 (CO1) and the internal transcribed spacer (ITS) barcodes (Nolan et al. 2007; Pagès et al. 2009; Stephan et al. 2009; Mathieu et al. 2011; Lehmann et al. 2012).

A species-specific real-time PCR test targeting the ITS-1 gene region of C. *imicola* sensu stricto significantly improved the surveillance of this species in France, as it can be used to detect target species in mixed samples (Cêtre-Sossah et al. 2008). Subsequently, a quantitative duplex real-time TaqMan PCR assay was developed to determine the relative abundance of *C. obsoletus* and *C. scoticus* in pooled samples (Mathieu et al. 2011). Around the same period, Wenk et al. (2012) designed 11 real-time PCR assays targeting the mitochondrial COI gene for the specific identification of both vector and non-vector *Culicoides* species, including *C. chiopterus, Culicoides* deltus Edwards, 1939, *C. dewulfi, Culicoides* grisescens Edwards, 1939, C. *imicola, Culicoides* lupicaris Downes & Kettle, 1952, *C. obsoletus, C. pulicaris, C. scoticus*, and *Culicoides* sp. However, cross-species reactivity was observed for *C. chiopterus, C. scoticus*, C. deltus, C. grisescens, and *C. obsoletus*. Dähn et al. (2025) later developed TaqMan quantitative real-time PCR assays targeting six members of the *Culicoides* subgenus Avaritia: *C. obsoletus* clades O1, O2, and O3, *C. scoticus* clade 1, *C. chiopterus*, and *C. dewulfi*. Although these assays were designed for use in both singleplex and multiplex PCR formats, the study highlighted cross-reactivity in the multiplex format, which was attributed to overlapping primer-probe interactions. Several *Culicoides* species and distinct haplotypes have been identified in the Palearctic region using a combination of morphological and molecular characterisation. The presence of different haplotypes poses challenges for species-specific assay design, as some may be inadvertently excluded from detection or may lead to cross-reactivity with existing assays when degenerate primers are used to account for sequence variation (Wenk et al. 2012). These issues are particularly problematic when analysing unsorted field specimens.

In Ireland, although morphological and molecular characterisation have facilitated the development and validation of the *Culicoides* species list (Collins et al. 2018; Isiye et al. 2025), no species-specific assays have been developed to target the important *Culicoides* species. The development of such assays would significantly advance the large-scale monitoring of these species, greatly supporting surveillance efforts and improving our understanding of their ecology and role as vectors, particularly of BTV and SBV. This study aimed to develop a high-throughput species-specific real-time PCR method targeting the internal transcribed spacer 2 (ITS2) region for the surveillance of important *Culicoides* species captured in light traps on farmlands in Ireland.

## Materials and methods

### Primer and probe design

A nucleotide BLAST (BLASTn) search was performed using the ITS sequences obtained from seven *Culicoides* biting midge species: *C. obsoletus, C. scoticus, C. chiopterus, C. dewulfi, C. pulicaris, C. punctatus*, and *Culicoides impunctatus* Goetghebuer, 1920, which were generated through DNA barcoding analysis by Isiye et al. (2025). The corresponding reference sequences from GenBank with the highest similarity to the Irish *Culicoides* sequences were selected, and multiple sequence alignment (MSA) was performed using the EMBL-EBI Clustal Omega program (Madeira et al., 2022). The alignment included the following sequences: *C. obsoletus* (MK893033), *C. scoticus* (MK893045), *C. chiopterus* (MK893002), *C. dewulfi* (MK893005), *C. pulicaris* (DQ371264), *C. punctatus* (DQ371247), and *C. impunctatus* (EU908207) (Gomulski et al., 2006; Mathieu et al., 2020). Conserved and variable regions were identified within the alignment, and the forward primer, reverse primer, and, for certain tests, a probe was designed based on interspecies variable regions to ensure test specificity, using Primer Express™ Software v3.0.1. Primer specificity was evaluated *in silico* by initially performing a Primer-BLAST analysis using the NCBI Primer Design Tool.

### Efficiency testing

The efficiency of the novel real-time PCR primers was evaluated using a 5-log dilution series of target species DNA obtained from a previous study by Isiye et al. (2025). Each reaction consisted of 5 µL of 2x FastStart Universal SYBR Green Mastermix (ROX) (<u>Sigma-Aldrich</u>), 0.4 µL of forward and reverse primer mix with a final concentration of 200 nM, 3.6 µL of molecular-grade water, and 1 µL of extracted DNA as the template. The negative control reactions included 1 µL of molecular-grade water instead of the DNA template. Reactions were prepared in a final volume of 10 µL and performed in triplicate using a QuantStudio™ 3 Real-Time PCR System (Applied Biosystems). The real-time PCR profile consisted of an initial incubation at 50 °C for 2 min, followed by denaturation at 95 °C for 10 min, and 40 cycles of 95 °C for 15 s and 60 °C for 1 min. Primer specificity and sensitivity were assessed using amplification and melt curve analyses. A standard curve was generated, and the primer efficiency (E) was calculated, with the recommended efficiency range being 90–110%. The curve was visualised using the ggplot2 package in R version 4.4.1 (R Core Team, 2025).

### Cross-species reactivity testing

To determine cross-species amplification, all primer sets were tested against 0.25 ng DNA of each non-target species, as described above. All assays were first tested as SYBR-based assays; however, for assays that amplified the target and non-target species, a probe was introduced to increase the specificity of the tests. Each probe-based reaction contained 5 µL TaqMan™ Fast Advanced Master Mix (Applied Biosystems), 0.2 µL of forward and reverse primers, probe mix with a final concentration of 250 nM, 3.8 µL of molecular-grade water, and 1 µL of extracted DNA. Negative controls were prepared by replacing the DNA template with 1 µL molecular-grade water.

### Application to mixed field trap samples

*Culicoides* specimens collected in June 2023 from 13 study sites were pooled into groups of individual midges, resulting in 13 pooled samples representing field trap collections. The pooled midge samples were crushed in extraction buffer using a pipette tip, DNA was extracted from each pool, and its concentration was measured using a NanoDrop™ 8000 spectrophotometer. Each extracted DNA sample was subjected to a 1:10 dilution and used as template DNA. All reactions were performed in triplicate for both SYBR-based and hydrolysis probe-based assays. The reaction conditions followed the respective protocols previously established during cross-species reactivity testing, and the average Ct values and melting temperatures (Tm) were recorded. To visualise the presence or absence of each species within the mixed samples, a dot plot was generated in R version 4.4.1 (R Core Team, 2025) using the ggplot2 package.

## Results

### Primer and probe design

Seven real-time PCR primer sets were designed targeting the *Culicoides* species: *C. pulicaris, C. scoticus, C. dewulfi, C. chiopterus, C. obsoletus, C. punctatus*, and *C. impunctatus* (Table 1).

**Table 1.**
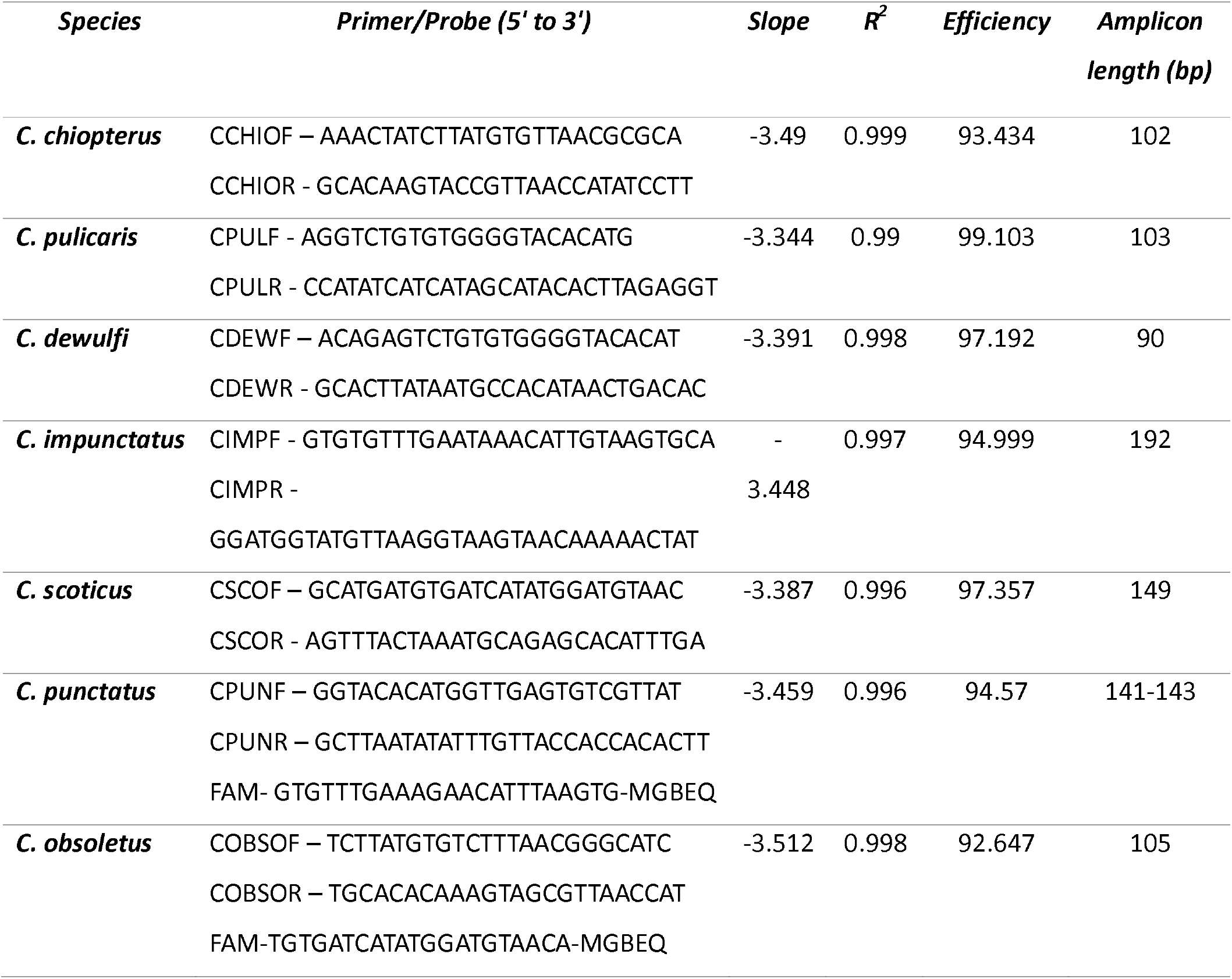
Primers and probes designed for the specific detection of Culicoides species, along with the standard curve results and efficiencies for the species-specific primer sets selected for each target species.

### Efficiency testing

When tested on the bench, the selected primer sets exhibited a well-defined dynamic range of amplification, producing five distinct amplification curves (Figures 1 and 2). Melt curve analysis further validated the specificity of amplification by generating distinct peaks at specific melting temperatures (Tm) (Supplemental Information). Additionally, standard curve analysis (Figures 1 and 2) confirmed that the primer efficiency remained within the optimal range of 90-100% (Table 1).

**Figure 1.**
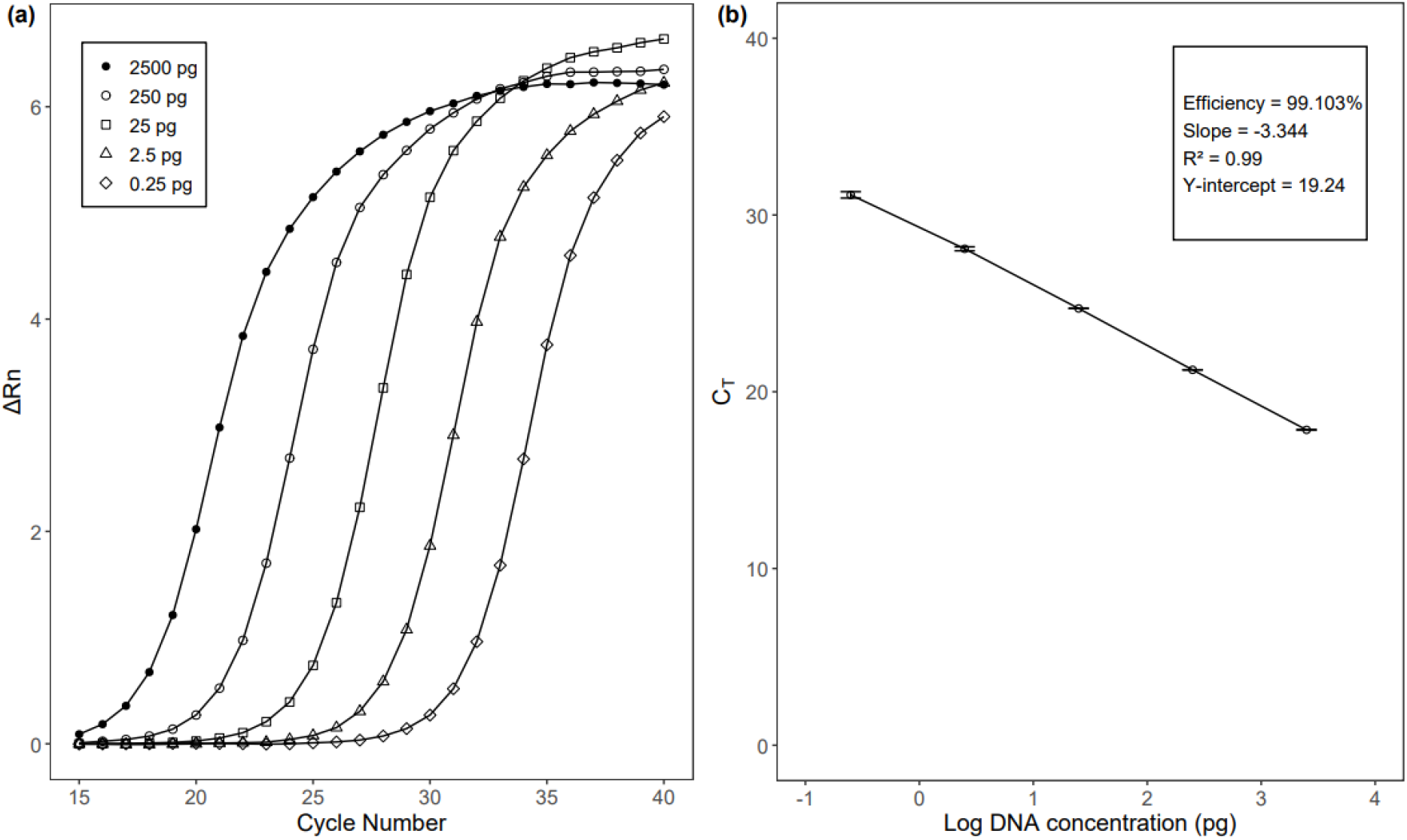
Amplification of Culicoides pulicaris target DNA using CPUL primers. (a) Amplification plot showing a 1:10 serial dilution series with corresponding amplification curves and (b) standard curve showing the linear regression of Ct values against DNA concentration, demonstrating assay efficiency and sensitivity.

**Figure 2.**
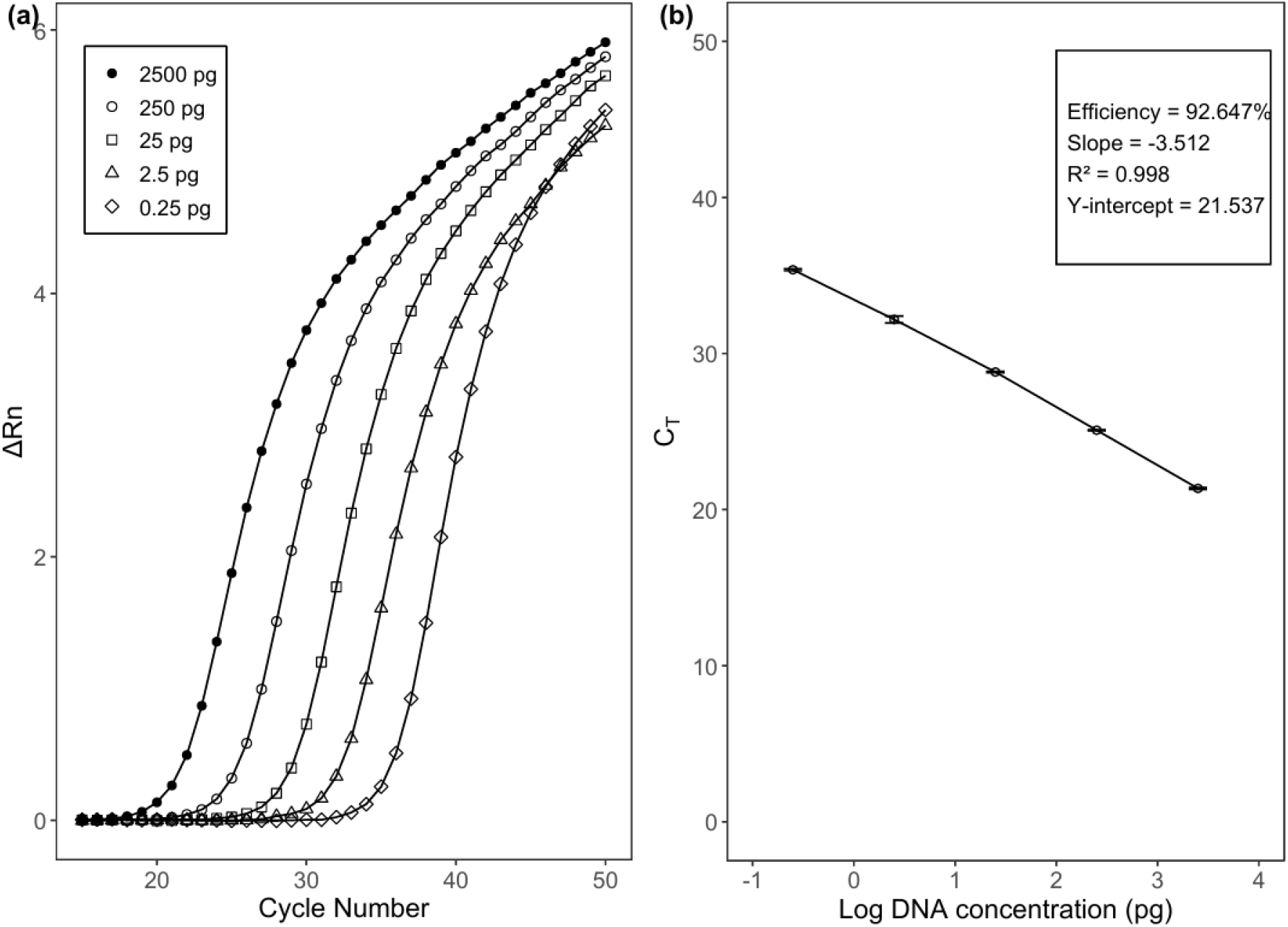
Amplification of Culicoides obsoletus target DNA using COBSO primers and a TaqMan® probe. (a) Amplification plot displaying a 1:10 serial dilution series with corresponding amplification curves and (b) Standard curve showing the linear regression of Ct values against DNA concentration, demonstrating assay efficiency and sensitivity.

### Cross-species reactivity testing

As shown in Figure 3, the assays effectively differentiated the target species while minimising non-specific amplification. Setting a cutoff Ct value of 34 ensured that false positives were not included, as any amplification beyond this value was not considered indicative of target detection. Furthermore, no cross-reactivity was detected in the assays developed for *C. pulicaris* (CPUL), *C. impunctatus* (CIMP), *C. scoticus* (CSCO), *C. dewulfi* (CDEW), and *C. chiopterus* (CCHIO). The designed SYBR-based assays lacked specificity for two species. These included *C. punctatus*, which was successfully amplified using the primers CPUNF and CPUNR at a Ct of 25. However, *C. impunctatus* was also amplified (at a Ct of 30), indicating some cross-reactivity. To enhance specificity, a probe was introduced, which improved the assay by amplifying *C. punctatus* at a Ct value of 23 while eliminating cross-reactivity with *C. impunctatus*. The second species was *C. obsoletus*, which was successfully amplified using COBSOF and COBSOR with a Ct of 21 and Tm of 71°C. However, non-target species, including *C. pulicaris, C. chiopterus, C. dewulfi, C. impunctatus, C. scoticus*, and *C. punctatus* were also amplified, though at Ct values >34. To improve specificity, a probe was introduced to eliminate non-target amplification, and the resulting Ct values are presented in Table 2. During cross-species reactivity testing of the *C. obsoletus* assay, occasional late amplification was observed at Ct values ranging from 35.6 to 38.6. These amplifications were inconsistent, occurring in only one or two of the three technical replicates, and were detected in some non-target species despite the inclusion of a species-specific probe, which reduced but did not fully eliminate these non-specific signals. No amplification was observed in any of the no-template control (NTC) assays. The average Ct and Tm values for all the assays are presented in Table 2.

**Table 2.** The average cycle threshold (Ct) and melting temperature (Tm) values for each species-specific assay using SYBR Green-based and probe-based real-time PCR. Note: Tm values are not reported for C. obsoletus and C. punctatus because probe-based assays do not utilize melting temperature analysis for target detection.

| **Assay name** | **Species** | **Average**<br>**Ct** | **Average Tm (°C)** | **Standard**<br>**deviation** |
| --- | --- | --- | --- | --- |
| *CCHIO* | *C. chiopterus* | 21 | 69 | 0.07 |
| *CPUL* | *C. pulicaris* | 21 | 70 | 0.09 |
| *CDEW* | *C. dewulfi* | 23 | 70 | 0.06 |
| *CIMP* | *C. impunctatus* | 27 | 68 | 0.48 |
| *CCSO* | *C. scoticus* | 26 | 69 | 0.07 |
| *CPUN* | *C. punctatus* | 23 | - | 0.21 |
| *COBSO* | *C. obsoletus* | 22 | - | 0.07 |

**Figure 3.**
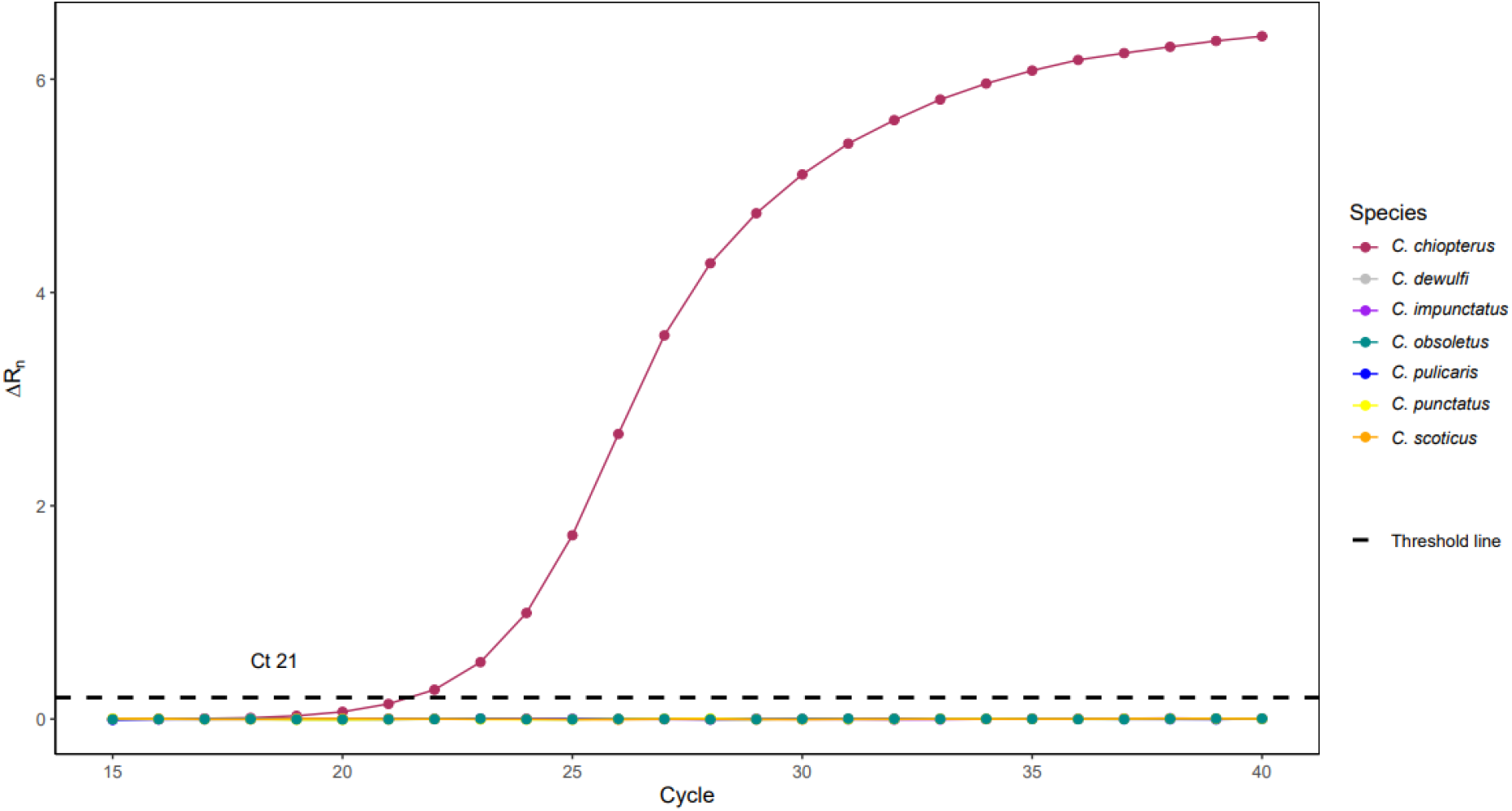
Amplification of C. chiopterus and non-target species using species-specific primers, ensuring selective amplification without cross-reactivity.

### Application to mixed field trap samples

A total of 840 *Culicoides* specimens, pooled into 13 samples from 13 distinct field sites (OVI 1 to OVI 13), were screened using the species-specific assays developed in this study (see Supplemental Information). DNA concentrations from the pooled extractions ranged from 3.0 to 27.3 ng/µL. Figure 4 displays a presence/absence dot plot indicating the detection of the target *Culicoides* species across the 13 pooled field samples. The corresponding cycle threshold (Ct) values for each assay are provided in the Supplemental Information.

**Figure 4.**
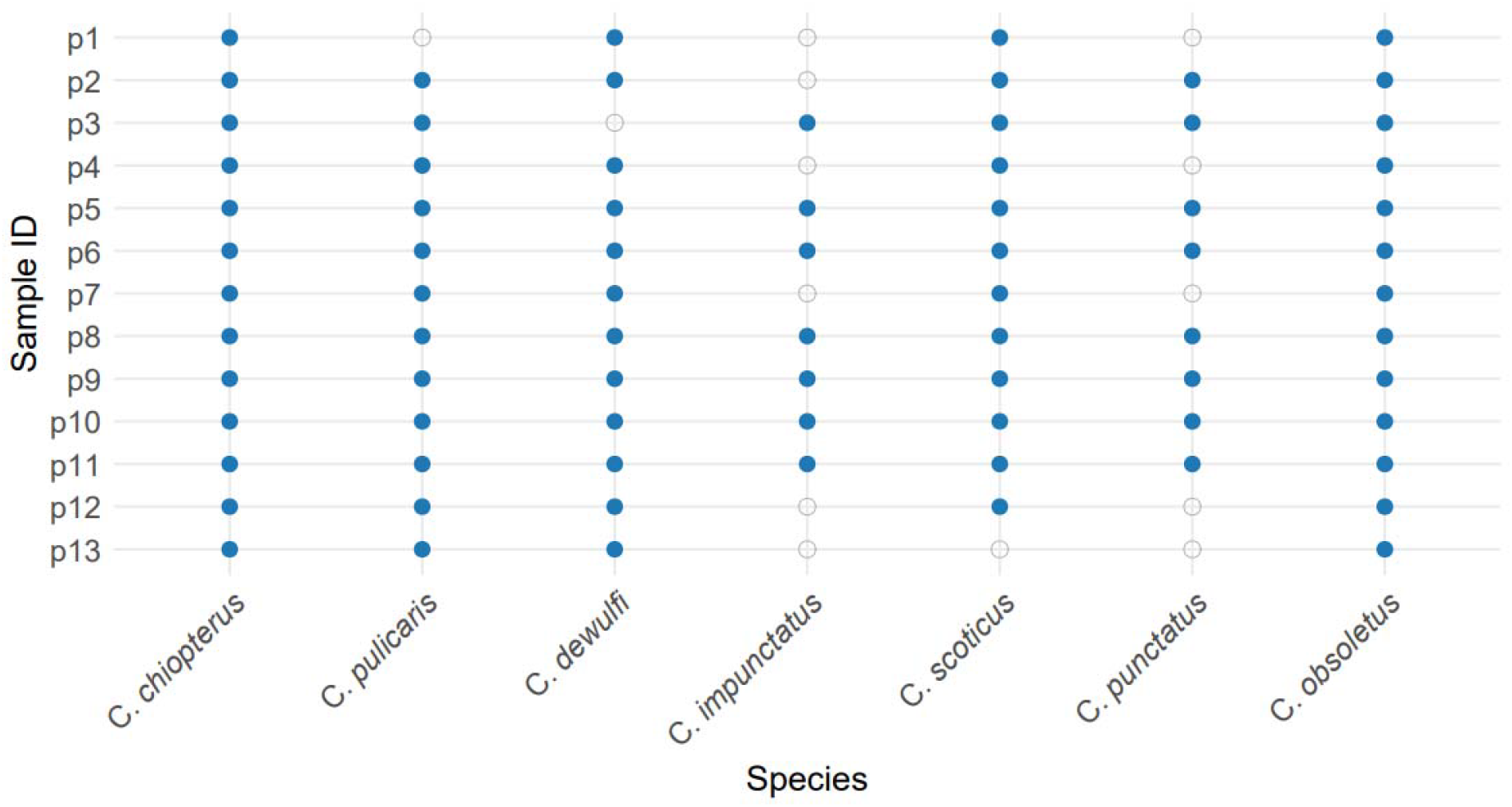
A dot plot displaying the presence or absence of Culicoides species across 13 pooled field samples, p1 to p13.

## Discussion

Advances in molecular techniques have significantly improved biodiversity monitoring, enhancing the efficiency of vector surveillance through accurate species discrimination and genetic comparative analyses across different geographic regions (Mignotte et al. 2021; Onyango et al. 2015; Tay et al. 2016). To aid in molecular surveillance, diagnostic tests have been developed by designing primers that target highly variable regions, ensuring high specificity and amplification efficiency. Tools such as Primer 3 and Primer Express™ Software are used to select primers, considering factors like melting temperature (Tm), primer length, GC content, and secondary structure (Rozen and Skaletsky, 2000). Species-specific real-time PCR assays are designed for high sensitivity, enabling the detection of a single *Culicoides* species within a mixed insect pool. Some assays have demonstrated the ability to detect a single C. *imicola* in a pool of 3,200 *Culicoides* or within 65 mg of material from a light trap (Cêtre-Sossah et al., 2004), while others can detect specific species in pools of up to 1,000 *Culicoides* (Wenk et al. 2012). The mitochondrial COI gene has been commonly used in the design of species-specific assays because of its low intraspecific variability, which accounts for haplotype diversity, and high interspecific variation (Aguilar-Vega et al. 2021; Balczun et al. 2009; Dallas et al. 2003; Pagès et al. 2009). However, for closely related or cryptic species, the more variable ITS region is often preferred (Dallas et al. 2003, Pages et al. 2009; Augot et al 2010; Curran et al. 2025). The ITS1 region was initially used to develop species-specific assays for C. *imicola* (Cêtre-Sossah et al. 2008), while later studies targeted the ITS2 region to design real-time PCR assays for *C. obsoletus, C. scoticus*, and *Culicoides montanus* Wirth & Blanton, 1969, demonstrating improved sensitivity compared to conventional assays (Monaco et al. 2010). The entire ITS1-5.8S-ITS2 region has also been used in quantitative duplex real-time PCR assays for detecting and quantifying *C. obsoletus* and *C. scoticus* (Mathieu et al. 2011). In this study, the ITS2 region was selected for primer design because it has been successfully used for the molecular characterisation of *Culicoides* species in Ireland, confirmed through both morphological and DNA barcoding of individual specimens (Isiye et al. 2025). This provided us with verified, high-quality sequences and ample genetically scored positive control samples for assay development and validation.

In this study, we developed five SYBR-based and two probe-based real-time PCR protocols for identifying *C. obsoletus, C. scoticus, C. chiopterus, C. dewulfi, C. pulicaris, C. punctatus*, and *C. impunctatus*, species commonly found across Ireland (McCarthy et al. 2009; Jess et al. 2018; Collins et al. 2018; Isiye et al. 2025). Primers were designed to account for haplotype diversity, while degenerate bases were avoided to minimise non-specific amplification and enhance PCR efficiency. Sequence alignments of the seven *Culicoides* species were used to design species-specific primers, following approaches similar to those described in previous studies (Nolan et al. 2007; Stephan et al. 2009; Mathieu et al. 2011). While primer design aimed to balance specificity and amplification efficiency (Dieffenbach et al. 1993), *in silico* predictions did not always align with results achieved at the bench; therefore, several primers were designed, and alternative primers were used in case of non-specificity. For species such as *C. impunctatus*, the reverse primer was designed to be longer than most typical real-time PCR primers to ensure adequate specificity owing to the lack of a potential variable region in the alignment. The primary aim was to design SYBR assays, as they are more economical than TaqMan® probes. While this was successfully achieved for five species (*C. pulicaris, C. scoticus, C. impunctatus, C. dewulfi*, and *C. chiopterus*), for *C. obsoletus* and *C. punctatus*, specificity was increased through the inclusion of TaqMan® probes.

All primers used in this study accounted for haplotype diversity within each *Culicoides* species, as demonstrated by the Primer-BLAST results. Specifically, CPUL matched eight *C. pulicaris* and five C. lupicaris haplotypes; CIMP matched two *C. impunctatus*; CDEW matched 20 *C. dewulfi*; CCHIO matched seven *C. chiopterus*; CPUN matched eight *C. punctatus*; and CSCO matched 14 *C. scoticus*. However, the COBSO primers matched a wider range of haplotypes, including 21 haplotypes of *C. obsoletus*, 15 of *C. montanus*, two of *Culicoides* sinanoensis Tokunga, 1937, and one of *C. scoticus* (Supplemental Information). Further analysis by sequence alignment confirmed that the COBSO probe successfully excluded most non-target species; however, it could likely bind to one haplotype each of *C. sinanoensis* (MK893047), *C. scoticus* (FN263316), and *C. montanus* (MK893025). Representative specimens for these three specific haplotypes were not available, preventing experimental validation of this limitation. The *C. sinanoensis* haplotype identified in Russia (Mathieu et al., 2020) belongs to the eastern Palearctic obsoletus complex. Primer-BLAST results detected four SNPs, three on the forward primer and one on the reverse primer, potentially increasing the Ct values if amplified. The *C. scoticus* haplotype, deposited by Kiehl et al. (2009), was collected in Germany (2007–2008) after the 2006 BTV outbreak. In their study, this *C. scoticus* haplotype was considered a *C. obsoletus* variant rather than a distinct species. However, recent studies have confirmed it as a distinct cryptic species (Mathieu et al. 2020; Mignotte et al. 2020). Finally, *C. montanus* is a rare species with a limited distribution in the Palearctic region (Delécolle et al. 2002; Mathieu et al. 2007). It has been observed in Morocco, areas with high land surface temperatures in Italy, and in small numbers in France, Norway, and Spain (Mathieu et al. 2012; Balenghien et al. 2018; Antoine et al. 2020; Aguilar-Vega et al. 2021). Morphologically, *C. montanus* is part of the obsoletus complex and is characterised by a deep sensory pit and hypertrophied trichoid sensilla (hair-like structures of different lengths) (Mathieu et al. 2011). Genetically, *C. montanus* and *C. obsoletus* form a monophyletic clade with a low genetic divergence between them (Garros et al., 2014; Mathieu et al. 2011). Despite efforts to differentiate between them using species-specific assays, complete discrimination between *C. obsoletus* and *C. montanus* remains elusive. Therefore, the accuracy of PCR assays is hindered by unresolved phylogenetic relationships and the high genetic similarity shared between the two species (Aguilar-Vega et al. 2021; Dähn et al. 2025). *C. montanus* has not been reported in Ireland, and its vector competence and public health significance in Europe are largely unstudied, so it was not prioritised in this study. However, primers designed by Monaco et al. (2010) for *C. montanus* based on the ITS2 region could be useful in regions where *C. montanus* and *C. obsoletus* occur sympatrically.

A Ct cut-off value of 34 was used in all assays to improve the test accuracy, ensuring good sensitivity and specificity (Greiner, 1995; Caraguel et al. 2011). This ensured that only reliable amplification signals were interpreted as positive. The cut-off was based on primer efficiency testing, balancing the detection of low DNA concentrations, and minimising false positives/negatives. For assays such as that developed for *C. obsoletus*, the application of this Ct cut-off value effectively excluded the high Ct signals observed during cross-species reactivity testing (Supplemental Information), which likely reflect low-level, non-specific amplification or background noise. Implementing this threshold also minimises the risk of false positives when the assay is applied to mixed-species field traps. Overall, the results indicate that cross-species amplification is minimal and unlikely to compromise the accuracy or reliability of the assay. When used to analyse trap samples, our real-time PCR assays can provide a semi-quantitative assessment of the number of target species in individual light trap catches, with lower Ct values indicating higher target DNA concentrations. Cêtre-Sossah et al. (2008) emphasised that detecting higher numbers of species, like C. *imicola*, is more critical for risk assessment than low numbers of individuals. In this study, Ct values were used to estimate the relative abundance of each *Culicoides* species within the mixed-species pools. Lower Ct values were interpreted as indicative of higher DNA concentrations and, therefore, a greater number of individuals of the corresponding species. This approach provides an approximate estimation of species prevalence in the pooled samples. However, it is important to consider that interspecific variations in body size, such as the relatively larger size of *C. pulicaris* compared to other species, may influence DNA yield and thus affect Ct-based abundance estimates.

In addition to facilitating the rapid identification of adult midges in trap samples, species-specific real-time PCR assays should also be able to detect larvae in environmental samples. Vector control efforts for disease prevention typically focus on managing breeding sites, a strategy that has been extensively applied to mosquito species. While different species of *Culicoides* larvae are thought to occupy different niches within the same site (Kettle and Lawson, 1952), little is known about their species-specific occupation in breeding sites. While conventional PCR has been used to identify larval samples (Yanase et al., 2013), real-time PCR is recommended for large-scale studies, as it is more high-throughput, rapid, and effective for identifying breeding sites in farming environments, which are usually linked to the spread of vector species and disease transmission. The assays can also be used for biodiversity monitoring by enabling precise detection of *Culicoides* species diversity across habitats, supporting fine-scale habitat association studies, and revealing ecological preferences and potential shifts in emergence patterns driven by climate change or land use changes. Overall, this molecular approach can be used to enhance the ecological understanding of *Culicoides* communities in the Palearctic region, supporting both conservation efforts and the risk management of vector-borne diseases.

Although this method is highly efficient, designing species-specific primers for all species that occur in Ireland was not possible because of the high number of species present. Therefore, this study focused on designing species-specific real-time PCR assays for the most common *Culicoides* species, as well as those most likely to drive BTV/SBV transmission in Ireland, to improve year-round monitoring of *Culicoides* biodiversity.

## Conclusion

The assays developed in this study successfully detected and discriminated all seven target *Culicoides* species, providing a valuable diagnostic tool for monitoring species composition in mixed field collections. These species-specific assays offer a high-throughput, rapid, cost-effective, and reliable method for tracking the distribution of *C. obsoletus, C. scoticus, C. chiopterus, C. dewulfi, C. pulicaris, C. punctatus*, and *C. impunctatus* adults in trap samples. Although their ability to detect *Culicoides* larvae in environmental samples has yet to be tested, this application would greatly aid targeted surveillance and control efforts. Overall, the PCR tools developed in this study can help improve our understanding of *Culicoides* biodiversity and vector-pathogen dynamics.

## Acknowledgements

This research was conducted as part of the Network of Insect Vectors (NetVec) Ireland project (https://www.ucd.ie/netvecireland/), funded by the Department of Agriculture, Food, and the Marine. The authors sincerely thank the NetVec Ireland team and the midge survey volunteers across Ireland for their efforts in specimen collection. The invaluable support and cooperation of these collaborators were instrumental in the successful completion of this study.

