## Supplemental Information for "Design and optimisation of rapid real-time PCR assays for the detection of key *Culicoides* species"

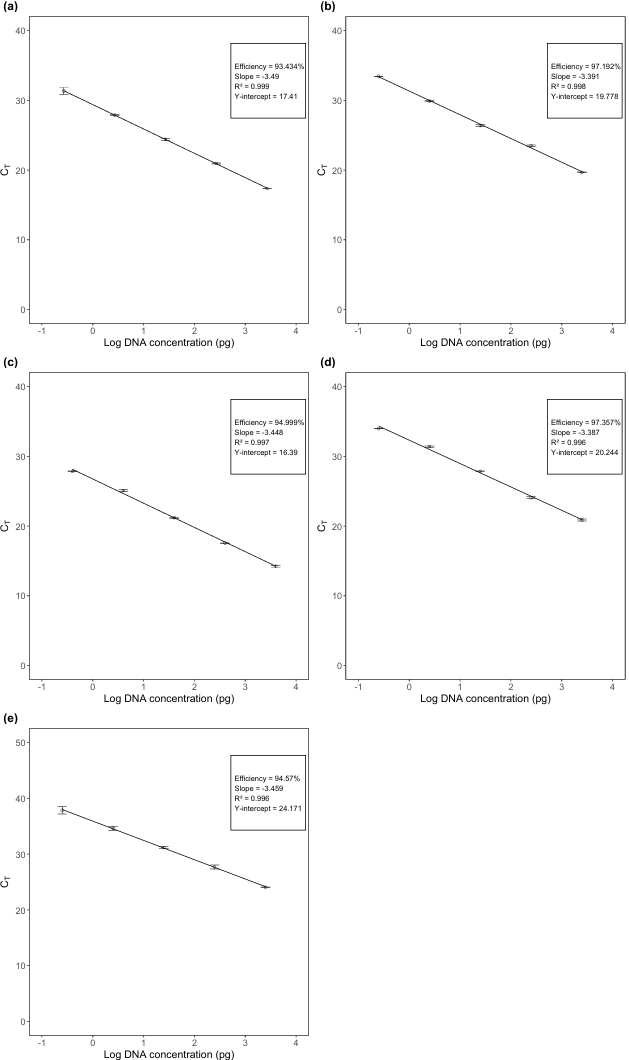

**Figure 5: Standard curves for (a) Culicoides chiopterus, (b) C. dewulfi, (c) C. impunctatus, (d) C. scoticus, and (e) C. punctatus, showing linear regression of Ct values against DNA concentration.**

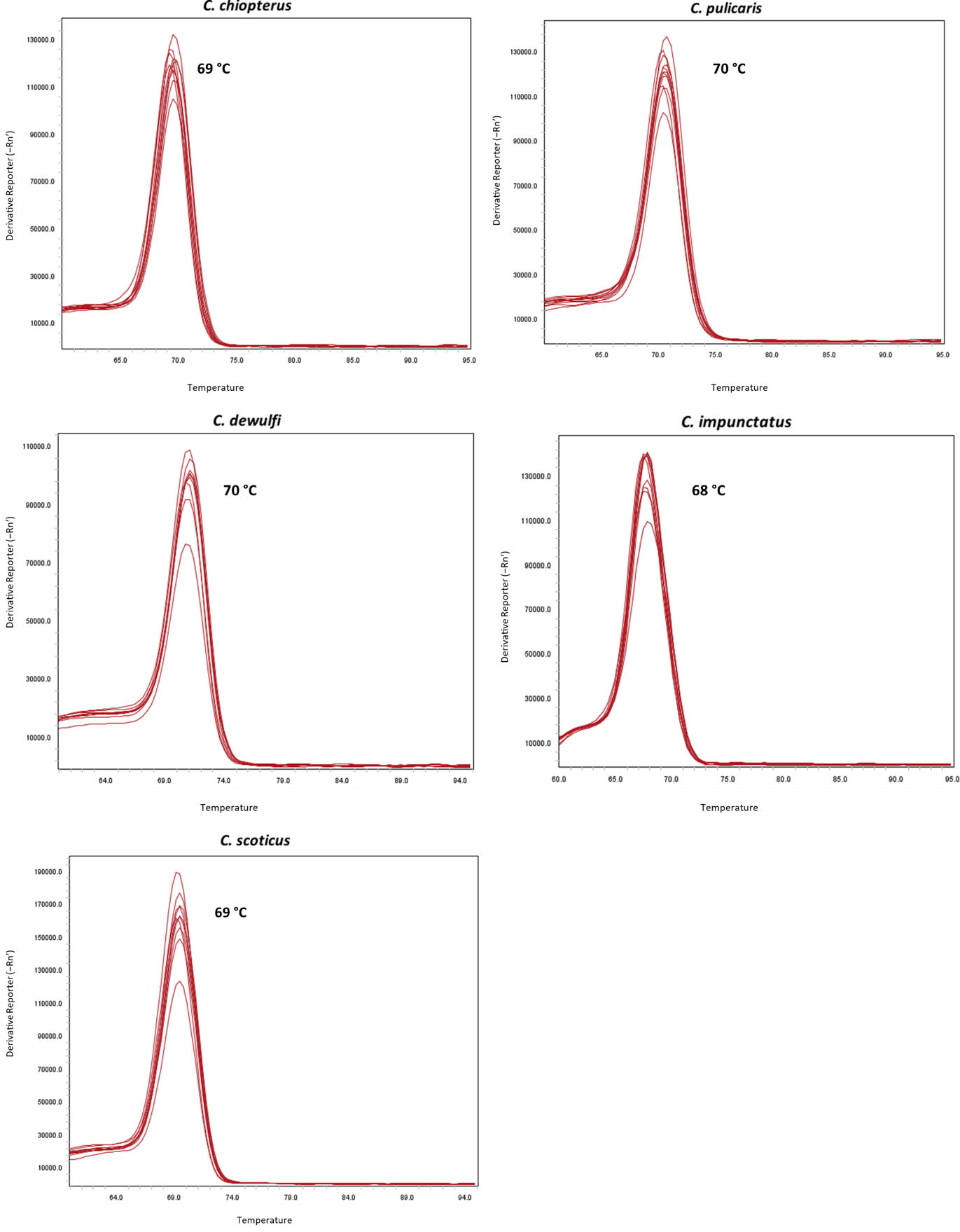

**Figure 6: Melt curve analysis for each species-specific SYBR-based test, showing distinct melting temperature (Tm) peaks corresponding to each Culicoides species target.**

***Table 3: Summary of 13 pooled DNA samples collected from 13 field sites, including DNA concentrations and corresponding Ct values for each species-specific assay. Results confirm the presence of individual Culicoides species within each pooled sample based on amplification in the respective tests.***

| ***Site ID*** | ***County*** | ***Sample ID*** | **No. of midges per pool** | ***Concentration ng/µL*** | ***C. chiopterus*** | ***C. pulicaris*** | ***C. dewulfi*** | ***C. impunctatus*** | ***C. scoticus*** | ***C. punctatus*** | ***C. obsoletus*** |
| --- | --- | --- | --- | --- | --- | --- | --- | --- | --- | --- | --- |
| OVI 1 | Dublin | *p1* | *21* | 3.035 | 21.7 | 0 | 21.9 | 0 | 23.1 | 0 | 25.8 |
| OVI 2 | Meath | *p2* | *60* | 22.12 | 16.3 | 18.2 | 16 | 0 | 21.7 | 27.2 | 20.8 |
| OVI 3 | Laois | *p3* | *32* | 1.975 | 32 | 21.4 | 0 | 26.5 | 29 | 28.1 | 27.6 |
| OVI 4 | Cork | *P4* | *92* | 14.93 | 19.8 | 32.7 | 16.6 | 0 | 24.5 | 0 | 20.1 |
| OVI 5 | Kerry | *p5* | *89* | 6.565 | 22.7 | 22 | 21 | 28.8 | 30 | 24.6 | 21.8 |
| OVI 6 | Carlow | *p6* | *88* | 15.04 | 21.7 | 21.3 | 22.3 | 26 | 33.7 | 24.5 | 19.6 |
| OVI 7 | Carlow | *p7* | *75* | 15.09 | 17.3 | 32.6 | 15.3 | 0 | 22.7 | 0 | 18.9 |
| OVI 8 | Galway | *p8* | *94* | 13.98 | 30.2 | 20.6 | 18.5 | 18.3 | 21.3 | 22 | 18.7 |
| OVI 9 | Clare | *p9* | *41* | 27.33 | 18.2 | 0 | 19.9 | 31.3 | 27.5 | 20.5 | 20.5 |
| OVI 10 | Limerick | *p10* | *96* | 13.79 | 18.7 | 30 | 16 | 24.9 | 21.2 | 33.3 | 20.9 |
| OVI 11 | Mayo | *p11* | *37* | 14.75 | 18.5 | 31 | 21.7 | 20.6 | 22.1 | 27.1 | 19.9 |
| OVI 12 | Waterford | *p12* | *22* | 5.209 | 32.4 | 33.5 | 17.5 | 0 | 20.3 | 0 | 21.8 |
| OVI 13 | Kilkenny | *p13* | *93* | 19.02 | 20.4 | 19.8 | 18.6 | 0 | 0 | 0 | 20.2 |

**Table 4: Results of in silico specificity testing for each species-specific primer, including predicted haplotypes and sequence alignments and sequences from non-target species with potential alignment to the COBSO probe.**

| **CCHIO** | **CPUL** | **CDEW** | **CIMP** |
| --- | --- | --- | --- |
| *C. chiopterus MK893002* | *C. pulicaris DQ371264* | *C. dewulfi MK893005* | *C. impunctatus EU908207* |
| *C. chiopterus MK893001* | *C. pulicaris DQ371266* | *C. dewulfi MK893003* | *C. impunctatus PQ859515* |
| *C. chiopterus PQ859511* | *C. pulicaris DQ371267* | *C. dewulfi PQ859514* |  |
| *C. chiopterus PQ859508* | *C. pulicaris DQ371268* | *C. dewulfi PQ859512* |  |
| *C. chiopterus EU908206* | *C. pulicaris DQ371265* | *C. dewulfi AY599831* |  |
| *C. chiopterus PQ859510* | *C. pulicaris PQ859518* | *C. dewulfi AY599830* |  |
| *C. chiopterus PQ859509* | *C. pulicaris PQ859519* | *C. dewulfi AY599828* |  |
|  | *C. pulicaris PQ859517* | *C. dewulfi AY599827* |  |
|  | *C. lupicaris DQ371255* | *C. dewulfi AY599826* |  |
|  | *C. lupicaris DQ371254* | *C. dewulfi AY599826* |  |
|  | *C. lupicaris DQ371252* | *C. dewulfi AY599824* |  |
|  | *C. lupicaris DQ371253* | *C. dewulfi AY599823* |  |
|  | *C. lupicaris DQ371256* | *C. dewulfi AY599822* |  |
|  |  | *C. dewulfi AY599821* |  |
|  |  | *C. dewulfi PQ859513* |  |
|  |  | *C. dewulfi AY599829* |  |
|  |  | *C. dewulfi AY599820* |  |
|  |  | *C. dewulfi AY599819* |  |
|  |  | *C. dewulfi AY599818* |  |
|  |  | *C. dewulfi MK893004* |  |

| **CSCO** | **CPUN** | **COBSO** | **COBSO + Probe** |
| --- | --- | --- | --- |
| *C. scoticus MK893045* | *C. punctatus PQ859520* | *C. obsoletus MK893037* | *C. obsoletus MK893037* |
| *C. scoticus MK893044* | *C. punctatus DQ371247* | *C. obsoletus MK893036* | *C. obsoletus MK893036* |
| *C. scoticus PQ859505* | *C. punctatus DQ371246* | *C. obsoletus MK893034* | *C. obsoletus MK893034* |
| *C. scoticus JF280793* | *C. punctatus DQ371248* | *C. obsoletus MK893033* | *C. obsoletus MK893033* |
| *C. scoticus AY599811* | *C. punctatus DQ371249* | *C. obsoletus MK893032* | *C. obsoletus MK893032* |
| *C. scoticus AY599808* | *C. punctatus PQ859521* | *C. obsoletus PQ849522* | *C. obsoletus PQ849522* |
| *C. scoticus AY599805* | *C. punctatus AB462275* | *C. obsoletus JF280792* | *C. obsoletus JF280792* |
| *C. scoticus AY599799* | *C. punctatus DQ371250* | *C. obsoletus AY599795* | *C. obsoletus AY599795* |
| *C. scoticus AY599798* |  | *C. obsoletus AY599794* | *C. obsoletus AY599794* |
| *C. scoticus AY599797* |  | *C. obsoletus AY599793* | *C. obsoletus AY599793* |
| *C. scoticus AY599796* |  | *C. obsoletus AY599792* | *C. obsoletus AY599792* |
| *C. scoticus AY599809* |  | *C. obsoletus AY599791* | *C. obsoletus AY599791* |
| *C. scoticus AY599801* |  | *C. obsoletus AY599790* | *C. obsoletus AY599790* |
| *C. scoticus AY599803* |  | *C. obsoletus AY599790* | *C. obsoletus AY599790* |
|  |  | *C. obsoletus AY599788* | *C. obsoletus AY599788* |
|  |  | *C. obsoletus AY599783* | *C. obsoletus AY599783* |
|  |  | *C. obsoletus AY599782* | *C. obsoletus AY599782* |
|  |  | *C. obsoletus AY599781* | *C. obsoletus AY599781* |
|  |  | *C. obsoletus AY599780* | *C. obsoletus AY599780* |
|  |  | *C. obsoletus FN263314* | *C. obsoletus FN263314* |
|  |  | *C. obsoletus AY599785* | *C. obsoletus AY599785* |
|  |  | *C. montanus MK893028* | *C. montanus MK893025* |
|  |  | *C. montanus MK893026* | **C. scoticus FN263316* |
|  |  | *C. montanus MK893025* | **C. sinanoensis MK893047* |
|  |  | *C. montanus AY599777* |  |
|  |  | *C. montanus AY599776* |  |
|  |  | *C. montanus AY599775* |  |
|  |  | *C. montanus AY599771* |  |
|  |  | *C. montanus AY599770* |  |
|  |  | *C. montanus AY599769* |  |
|  |  | *C. montanus AY599779* |  |
|  |  | *C. montanus MK893027* |  |
|  |  | *C. montanus AY599774* |  |
|  |  | *C. montanus AY599773* |  |
|  |  | *C. montanus AY599772* |  |
|  |  | *C. montanus AY599778* |  |
|  |  | *C. scoticus FN263316* |  |
|  |  | *C. sinanoensis MK893046* |  |
|  |  | *C. sinanoensis MK893047* |  |
